# Single-neuron and population measures of neuronal activity in working memory tasks

**DOI:** 10.1101/2023.06.18.545508

**Authors:** Rana Mozumder, Christos Constantinidis

## Abstract

Information represented in working memory is reflected in the firing rate of neurons in the prefrontal cortex and brain areas connected to it. In recent years, there has been an increased realization that population measures capture more accurately neural correlates of cognitive functions. We examined how single neuron firing in the prefrontal and posterior parietal cortex of two male monkeys compared with population measures in spatial working memory tasks. Persistent activity was observed in the dorsolateral prefrontal and posterior parietal cortex and firing rate predicted working memory behavior, particularly in the prefrontal cortex. These findings had equivalents in population measures, including trajectories in state space that became less separated in error trials. We additionally observed rotations of the stimulus space for different task conditions, which was not obvious in firing rate measures. These results suggest that population measures provide a richer view of how neuronal activity is associated with behavior, however, largely confirm that persistent activity is the core phenomenon that maintains visual-spatial information in working memory.

**NEW & NOTEWORTHY:** Recordings from large numbers of neurons led to a re-evaluation of neural correlates of cognitive functions, which traditionally were defined based on responses of single neurons, or averages of firing rates. Analysis of neuronal recordings from the dorsolateral prefrontal and posterior parietal cortex revealed that properties of neuronal firing captured in classical studies of persistent activity can account for population representations, though some population characteristics did not have clear correlates in single neuron activity.

## INTRODUCTION

The ability to maintain and manipulate information in memory over a period of seconds is commonly referred to as working memory (1). Early neurophysiological experiments in non-human primates identified neurons that not only respond to sensory stimuli but remain active during a period after the stimuli were no longer present; this “persistent activity” provided a neural correlate of working memory (2). Persistent activity has since been widely replicated in a variety of working memory tasks (3), including in human recordings (4). Computational models typically simulate persistent activity by networks of neurons with recurrent connections between units similarly tuned for spatial location (5, 6). Activation in the network behaves as a continuous attractor (7), with initial activation by the stimulus appearance generating a bump (peak) of activity in the network, which is maintained in the delay period, but may drift randomly. The peak of the bump at the end of the delay period determines the recalled location (8). Drifts in neuronal activity from trial-to-trial account for corresponding deviations of behavior.

While analysis of this sort has placed emphasis on activity of individual neurons, or averages across multiple neurons that generate persistent activity, in recent years, it has been recognized that populations of neurons encode information that may not be readily visible in single neuron responses (9, 10). More generally, exploring the dynamics of population responses with dimensionality reduction methods have revealed latent variables that more accurately capture information about stimuli and task than individual responses (11, 12).

We were therefore motivated to compare aspects of single neuron and population activity that predict behavior. We analyzed recordings obtained from the dorsolateral prefrontal and posterior parietal cortex of monkeys as they performed working memory tasks, for which strong convincing evidence exists that single neuron activity is predictive of behavior. We examined neurons with persistent activity with conventional methods and performed dimensionality reduction methods to obtain population measures.

## METHODS

Data were obtained from two male rhesus monkeys (*Macaca mulatta*), weighing 7–9 kg, as previously documented (13), and were analyzed in this study. Monkeys were either single-housed or pair-housed in communal rooms with sensory interactions with other monkeys. All experimental procedures followed guidelines set by the U.S. Public Health Service Policy on Humane Care and Use of Laboratory Animals and the National Research Council’s Guide for the Care and Use of Laboratory Animals and were reviewed and approved by the Wake Forest University Institutional Animal Care and Use Committee.

### Experimental setup

Monkeys sat with their heads fixed in a primate chair while viewing a monitor positioned 60 cm away from their eyes with dim ambient illumination and were required to fixate on a 0.2° white square appearing in the center of the screen. During each trial and in order to receive a liquid reward (typically fruit juice) the animals maintained fixation on the square while visual stimuli were presented either at a peripheral location or over the fovea. Any break of fixation immediately terminated the trial, and no reward was given. Eye position was monitored throughout the trial using a non-invasive, infrared eye position scanning system (model RK-716; ISCAN, Burlington, MA). The system achieved a < 0.3° resolution around the center of vision. Eye position was sampled at 240 Hz, digitized and recorded. The visual stimulus display, monitoring of eye position, and synchronization of stimuli with neurophysiological data were performed with in-house software implemented on the MATLAB environment (MathWorks, Natick, MA), utilizing the Psychophysics Toolbox (14).

### Behavioral task

The task involved two stimuli appearing in sequence, requiring the monkey to make an eye movement to the remembered location of either the first or the second stimulus depending on the color of fixation point (Fig. 1). The animals fixated at a 0.2° fixation target located at the center of the monitor for 1 second, and then two white squares (1.5° in size) were displayed sequentially for 0.5 s, with a 1.5 s intervening delay period between them. The first stimulus (cue) was displayed pseudo-randomly at one of two locations, always drawn from eight possible locations along a circular ring of 12° eccentricity. The second stimulus location was either the same as the first, or differed by an angle of 45°, 90°, or 180°; a final condition involved no second stimulus presentation. After a second delay period of 1.5s, the monkeys were required to saccade to the location of the first stimulus if the fixation point was white in color (remember-first condition), and to the location of the second stimulus if the fixation point was blue (remember-second condition). To minimize the uncertainty about the stimulus to be remembered, the remember-first and remember-second conditions were presented in blocks of trials. The animal was required to perform 10 correct trials of the remember-first task, involving presentation of the two stimuli at each possible combination used in the block. The monkeys were rewarded with fruit juice after making a correct saccade. When running these sessions, the monkey was typically tested first with a single-stimulus Oculomotor Delayed Response task (15) to map the receptive field of neurons recorded, so as to guide selection of the location of stimuli.

**Figure 1.**
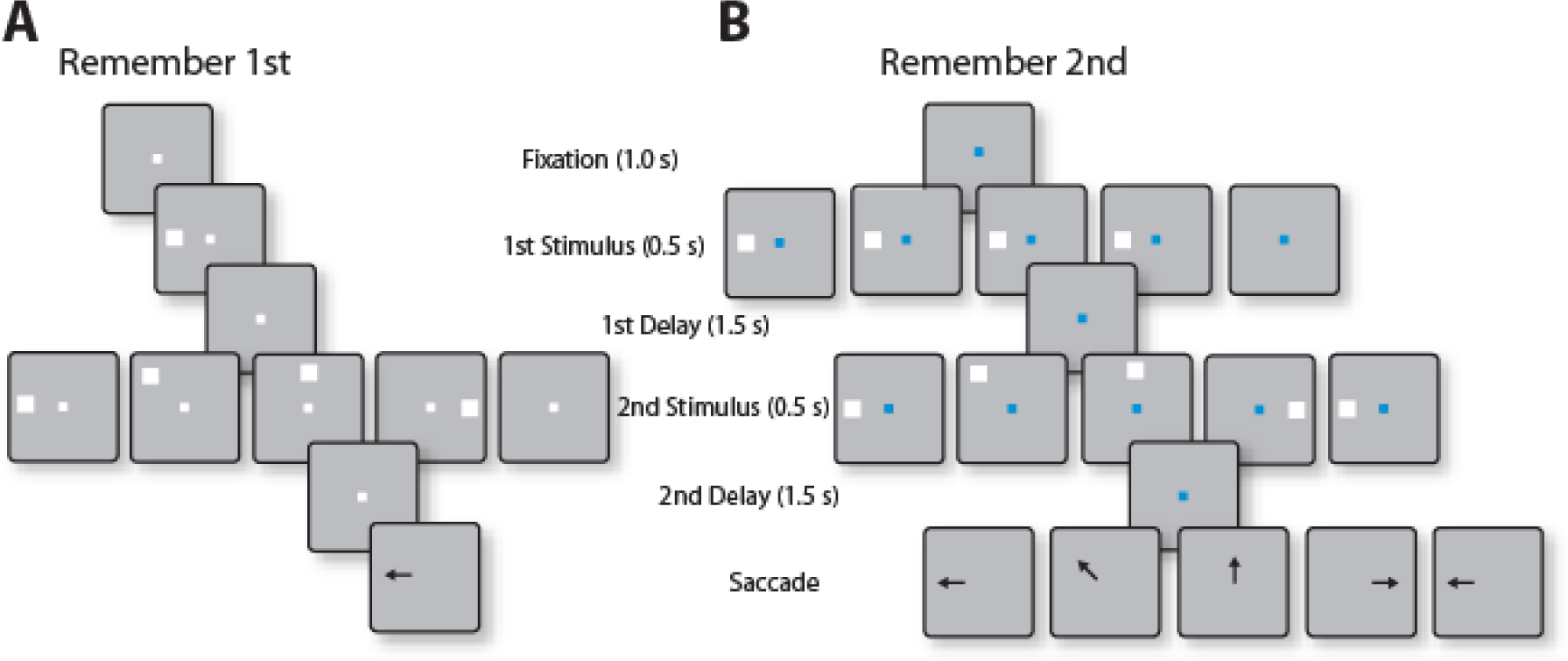
Successive frames show the sequence of stimulus presentations in the Remember First-Remember Second task. (A) In the remember first task, the fixation point was white, and the monkey had to remember the position of first stimulus (cue) and make a saccade to the position at the end of the trial while ignoring the second stimulus (distractor). Here the first stimulus could appear at either in the receptive field of the neuron (shown as the left location) or in the diametric position (not shown here), and depending on the position of the first stimulus, the distractor could appear at five different positions: at the same position as cue, at a location offset by 45, 90, 180 degrees, or no distractor at all (null condition). (B) In the remember second task, the monkey had to remember and make a saccade towards the second stimulus ignoring the first stimulus (distractor in this case). The cue could appear at the preferred or diametric location of the neuron followed by a second stimulus similar as before. In this case, the null condition involved no stimulus presentation in the first interval.

### Surgery and neurophysiology

Two 20 mm diameter recording cylinders were implanted over the dorsolateral prefrontal cortex and the posterior parietal cortex of the same hemisphere in each monkey (Fig. 2). Extracellular activity of single units were recorded using arrays of 2–8 microelectrodes in each cylinder, either with glass-coated, tungsten microelectrodes (250 μm diameter, impedance of 1 MΩ at 1 kHz, Alpha-Omega Engineering, Nazareth, Israel) or epoxylite-coated tungsten microelectrodes (125 or 250 μm diameter, impedance of 4 MΩ at 1 KHz, FHC Bowdoin, ME). Electrodes were advanced individually into the cortex with a Microdrive system (EPS drive, Alpha-Omega Engineering, Nazareth, Israel). The electrical signal from each electrode was amplified, band-pass filtered between 500 Hz and 8 kHz, and recorded with a modular data acquisition system at 25 μs resolution (APM system, FHC, Bowdoin, ME). The anatomical location of electrode penetration was determined based on MR imaging of the brain. Neuronal data were collected from areas 8a and 46 of dlPFC including both banks of the principal sulcus, and areas 7a and lateral intraparietal area (LIP) of the PPC in the lateral bank of the intraparietal sulcus.

**Figure 2.**
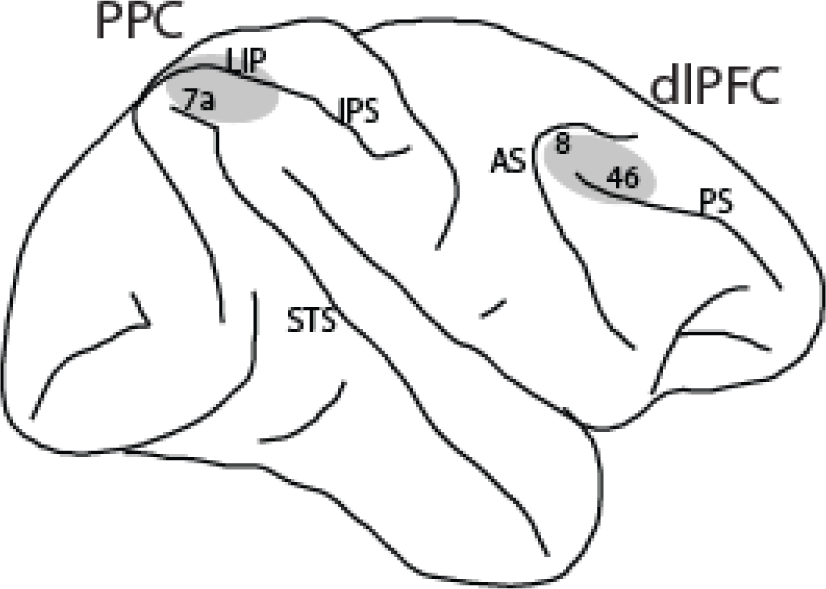
A schematic diagram of the monkey brain. Recordings were performed from the highlighted areas. In posterior parietal cortex (PPC), recordings were done from area 7a and lateral intraparietal area (LIP). In dorsolateral prefrontal cortex (dlPFC), recordings sampled area 8 and 46. IPS, intraparietal sulcus; STS, superior temporal sulcus; AS, arcuate sulcus; PS, principal sulcus.

### Neural data processing

A semi-automated cluster analysis relied on the KlustaKwik algorithm, which applied principal component analysis of the waveforms to sort recorded spike waveforms into separate units. To ensure a stable firing rate in the analyzed recordings, we identified recordings in which a significant effect of trial sequence was evident at the baseline firing rate (ANOVA, p < 0.05), e.g., due to a neuron disappearing or appearing during a run, as we were collecting data from multiple electrodes. Data from these sessions were truncated so that analysis was only performed on a range of trials with stable firing rate. Less than 10% of neurons were corrected in this way.

### Firing rate analysis

Data analysis was implemented with the MATLAB computational environment (MathWorks, Natick, MA). Neuron analysis involved, initially, determining the mean firing rate of each neuron in each trial epoch: 1 s of fixation; 0.5 s of cue presentation; 1.5 s of first delay period; 0.5 s of sample presentation; and 1.5 s of second delay period. Next, we compared responses of each neuron in the 1 s baseline, fixation period with the cue presentation period and the delay period following it. Any neuron that had significantly greater responses during the first delay period was identified as a task-responsive neuron (one-tailed paired t-test; p < 0.05).

In order to quantify the trial-to-trial association between neuronal activity and behavior, we analyzed trials that resulted in correct choices and incorrect choices using Receiver Operating Characteristic (ROC) analysis (16, 17). Firing rates of trials involving the same sequences of stimuli were pooled separately for correct and error outcomes and an ROC curve was computed from these two distributions of firing rates. The area under the ROC curve thus constructed is referred to in the literature as “choice probability” and represents a measure of correlation between the behavioral choice and neuronal activity. As we defined it for our analysis, a value of 1 indicates that higher firing rates of the neuron predict perfectly the behavioral choices of the subject; a value of 0.5 indicates no correlation between the two; a value of 0 indicates that lower firing rates of a neuron predict the behavioral choices. Time-resolved choice probabilities were computed from the spikes in 500 ms time windows, stepped by 50 ms intervals. This analysis was done separately for the preferred and non-preferred cue conditions.

### Dimensionality reduction

We applied principal components analysis (PCA) to analyze the neural population activity during the Remember-first and Remember-Second tasks. PCA was performed on the mean firing rate of neurons, for data collected from both PFC and PPC. Neurons from each brain region were pooled and data were collected for *L* = 10 different conditions in the Remember-First task. Each task condition was defined by the combination of the first and second stimulus locations: 5 conditions involved cue presentation at the neuron’s preferred location, and 5 at its diametric location; each condition was then followed by a different second stimulus location (as in Fig. 1A). The Remember-Second trials involved another 10 task conditions, structured in a similar fashion (as in Fig. 1B). For correct trajectories, we selected neurons with at least 6 correct trials per task condition. Then, for each neuron, the average firing rate of the Remember-first trials (and Remember-Second trials) during a given time point formed a column entry (10×1 vector) for the population activity matrix *A*_1_ (*A*_2_ for Remember-Second trials). During each time point, we defined the corresponding activity matrix *A*_1_ (*A*_2_) for each prefrontal region as an 10 x *N* matrix, where 10 is the number of task conditions and *N* is the number of neurons. We then aligned the activity matrices (*A*_1_ and *A*_2_) into one single matrix *B*, which is a 20 x *N* matrix with the first ten rows containing the activities in Remember-first task and the rest containing for Remember-Second. Then we normalized *B* by subtracting the mean across each column to guarantee the matrix was zero-centered. The PCA was applied on the centered data using singular value decomposition (SVD). We visualized data in the space defined by the first two principal components.

To create population activity trajectories, we used the binned firing rates of each neuron for each task condition, which comprised a matrix of *N* x 20 x *Number of bins*. For individual trajectories, we selected the respective task condition (matrix of *N* x *Number of bins* size) and multiplied with the inverse of PC1 and PC2 which were of *N*x 1 size each giving us two PC components of 1 x *Number of bins* size. Then we plotted them with respect to time.

For PCA analysis of error trials, we collapsed the 10 conditions of the Remember-First (and Remember-Second) task into two: those with the cue appearing at the neuron’s preferred location and those in the opposite. Then, we used binned firing rates as described, above to produce correct and error trajectories. To measure the distance between correct and error trajectories, we projected the trajectories to the state-space defined by the first two components and determined the average positions of the projected points across the delay period. In this way we obtained a point for each of four groups as described above and measured Euclidean distances between the points. corresponding to the best cue and diametric cue conditions. To make the method robust, we repeated the analysis by choosing 75% of the selected neurons and measuring average distance each time as described earlier. We repeated this analysis for 100 times and plotted the distribution of average distances.

## RESULTS

Neuronal data were analyzed from two monkeys trained to perform the Remember First – Remember Second task (Fig. 1). The task requires the monkeys to observe two stimuli presented in sequence, with intervening delay periods between them and to make an eye movement to the remembered location of either the first or the second stimulus depending on the color of the fixation point (white, or blue, respectively). Extracellular neurophysiological recordings were collected from areas 8a and 46 the dorsolateral prefrontal cortex and areas 7a and LIP of the posterior parietal cortex (Fig. 2). We identified neurons with significantly elevated responses during the first delay period of the task, as these were defined previously (13). We focused on data recorded from the randomized variant of the task, where the second stimulus could appear at a variable location relative to the first as illustrated in Fig. 1. A total of 205 neurons from dlPFC and 241 from PPC were recorded with this stimulus set. Of those, 75 neurons exhibited persistent activity in the delay period after the first stimulus presentation in the dlPFC (47 and 28, from the monkeys GR and HE, respectively) and 85 such neurons did so in the PPC (21 and 64, from monkeys GR and HE, respectively).

### Single neuron activity in correct and error trials

We examined activity of neurons that generated persistent activity in correct and error trials of the Remember First task. When the first stimulus, which the monkey needed to remember, appeared in each neuron’s preferred location, firing rate of prefrontal neurons was generally higher in correct compared to error trials (Fig. 3A). Mean firing rate averaged across the entire first delay period was significantly higher in correct than error trials (two tailed paired t-test, *t_176_*=2.93, p=0.004). In other words, trials in which persistent activity was diminished were more likely to result in errors. This evidence is consistent with prior results that have linked persistent, delay period activity with behavior in working memory tasks (15, 18, 19), now demonstrated in the context of a new task.

**Figure 3.**
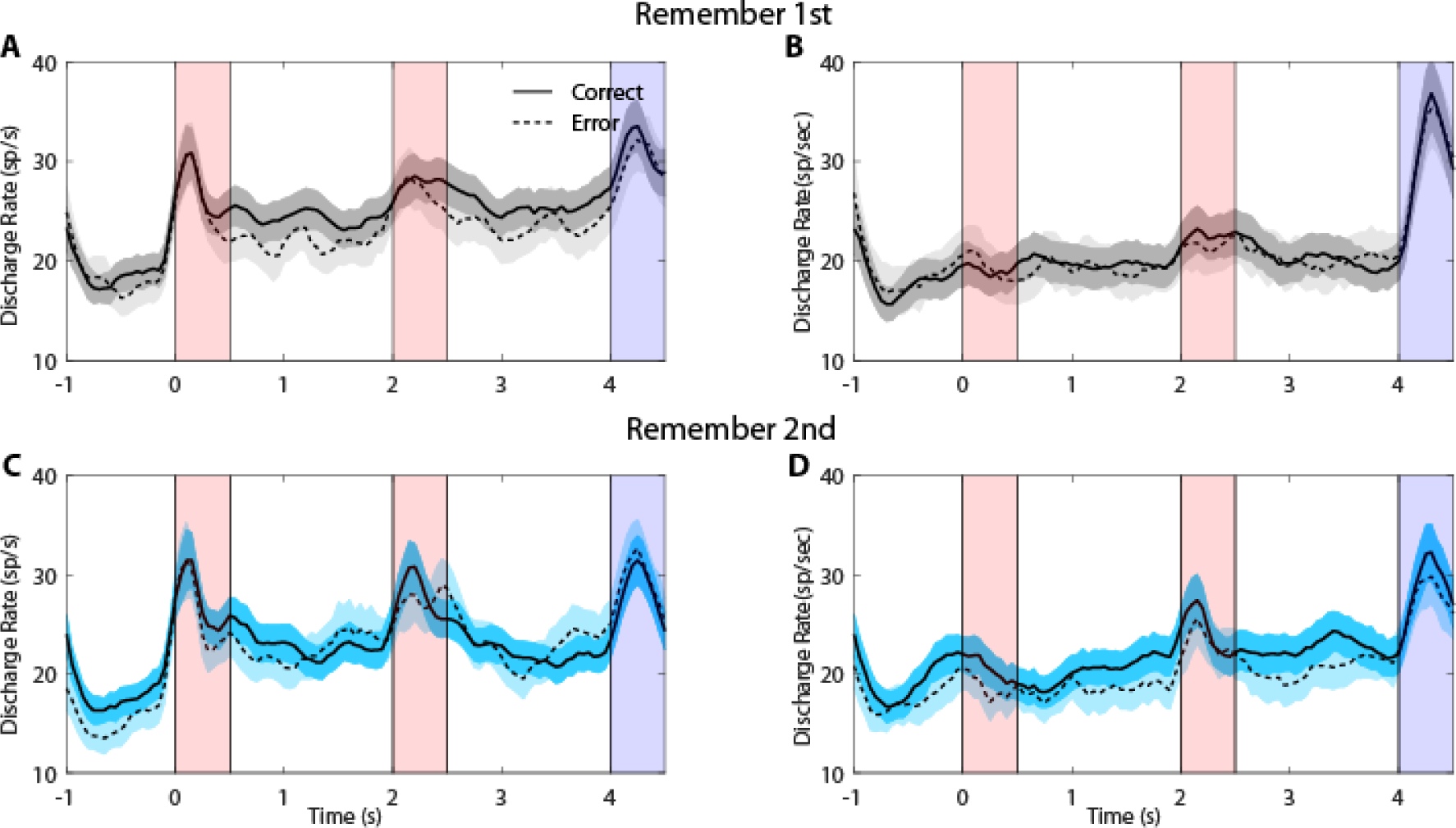
(A) Averaged Peristimulus Time Histogram (PSTH) of neuronal spike discharges from the PFC neurons that showed significantly elevated cue delay activity (n=75) in the remember first task when the cue appeared in the preferred location of the neurons. The solid line indicates mean activity of neurons in correct trials while the dashed line represents the averaged error trials activity. Shaded areas indicate standard error of mean (SEM). (B) As in A, for the remember first task when the cue appeared in the diametric position. (C) As in A, for remember second task when the cue appeared at the preferred location. (D) As in C, for remember second task when the cue appeared at the diametric location.

This difference in firing rate was not a general effect but depended on task conditions and differed between neuronal populations. The difference between correct and error trials was much more diminished following presentation of a stimulus in a neuron’s non-preferred location (Fig. 3B) and it did not reach statistically significance (two tailed paired t-test, *t_124_*=0.462, p=0.6). On the other hand, a trend in the opposite direction was evident in the Remember Second task, for a stimulus appearing in the neuron’s preferred location (Fig. 3C). When the monkey had to ignore this first stimulus, higher than average firing rate at the end of the first delay period was more likely to lead to an error. Finally, error trials involving appearance of a stimulus in a non-preferred location in the Remember Second task were characterized by lower firing rate, though this difference did not reach statistical significance (two tailed paired t-test, *t_149_*=1.78, p=0.078).

Importantly, the reduced firing rate in error trials involving presentation of the cue in the neuron’s receptive field was specific for neurons with persistent activity. Analysis of neurons that did not exhibit significantly elevated activity in the first delay period revealed generally higher firing rates in error than correct trials, across all conditions (Fig. 4). In other words, elevated activity by neurons that are ordinarily not active during the maintenance of working memory were more likely to lead to errors.

**Figure 4.**
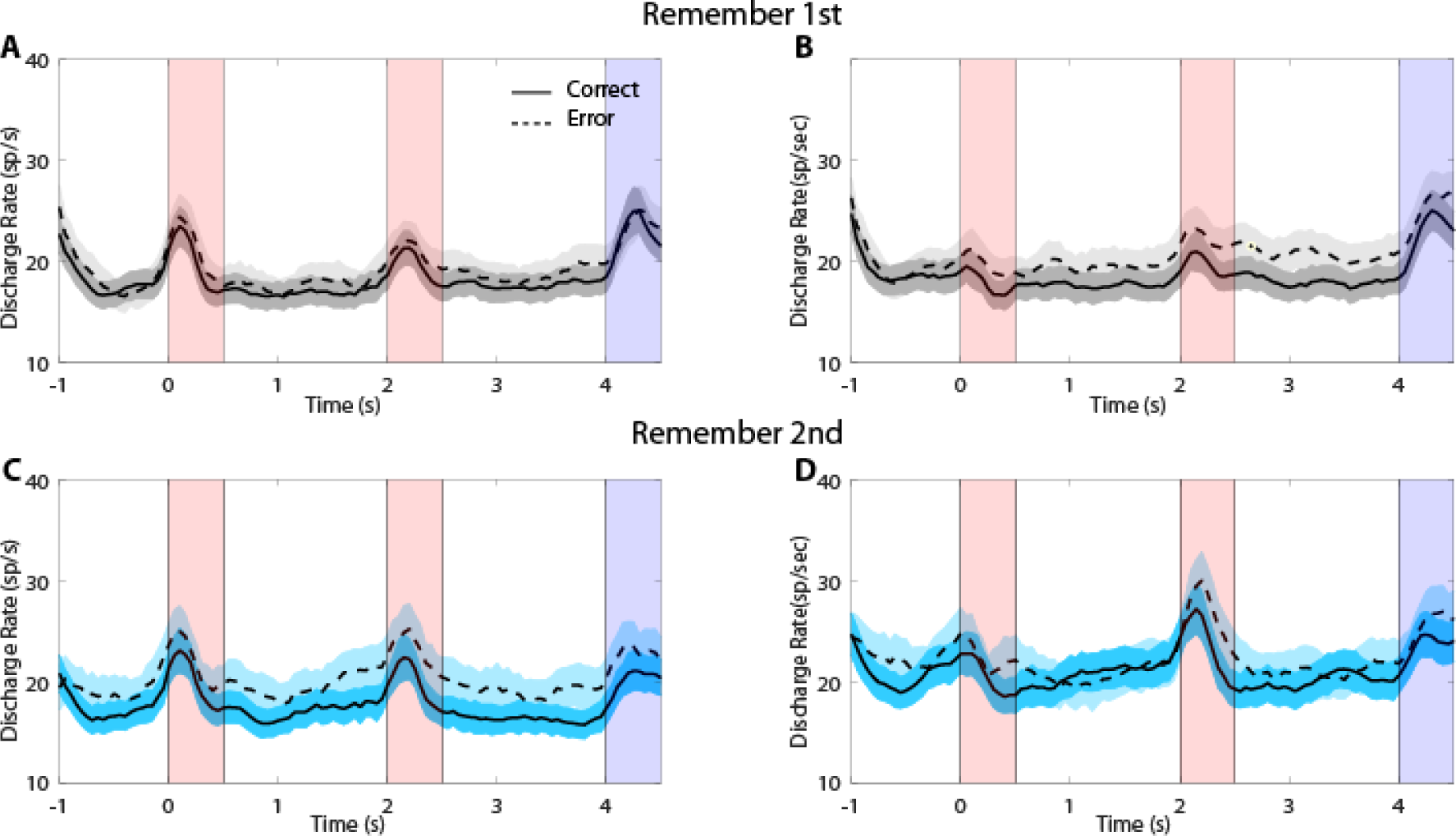
(A) Averaged PSTH of neuronal spike discharges from the PFC neurons that did not show significantly elevated cue delay activity (n=130) in the Remember-First task when the cue appeared in the preferred location of the neurons. Conventions are the same as in Fig. 3. (B) As in A, for the remember first task when the cue appeared in the diametric position. (C) As in A, for the remember second task when the cue appeared at the preferred location. (D) As in C, for the remember second task when the cue appeared at the diametric location.

Difference between correct and error trials was generally diminished among neurons that generated persistent activity in the posterior parietal cortex (Fig. 5). The firing rate for correct and error trials for the neurons’ best location in Fig. 5A did not reach statistical significance (two tailed paired t-test, *t_176_*=0.01, p=0.9). A number of previous studies have suggested that activity of posterior parietal neurons is less able to resist the effect of distractors (20) and firing rate differences have less influence on behavior.

**Figure 5.**
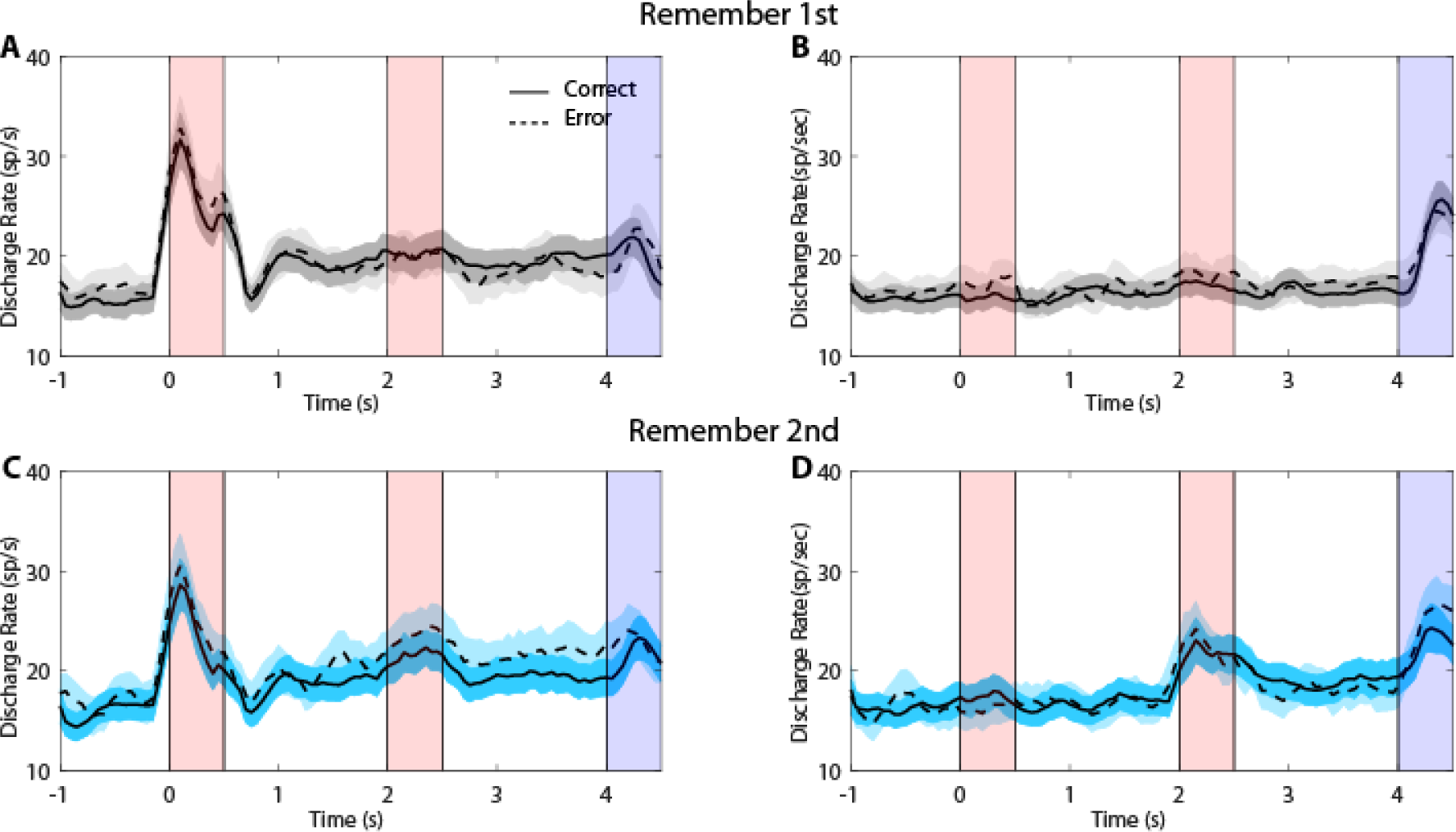
(A) Averaged PSTH of neuronal spike discharges from the PPC neurons that showed significantly elevated cue delay activity (n=85) in the remember first task when the cue appeared in the preferred location of the neurons. Conventions are the same in as in Fig. 3. (B) As in A, for the remember first task when the cue appeared in the diametric position. (C) As in A, for remember second task when the cue appeared at the preferred location. (D) As in C, for remember second task when the cue appeared at the diametric location.

Based on these results, we calculated “choice probability” for neurons that generated persistent activity, based on the area under the ROC curve defined by correct and error trials (Fig. 6), which is the conventional method of determining the influence of firing rate on behavior (17). Results were consistent with the analysis of firing rate, as presented above. Choice probability exceeded chance values (>0.5) in the delay interval following presentation of the cue in the neurons’ preferred location for the Remember-First task (Fig. 6, Left). Similarly, choice probability exceeded chance values in the delay interval following presentation of the cue in the neuron’s non-preferred location (Fig. 6, Right).

**Figure 6.**
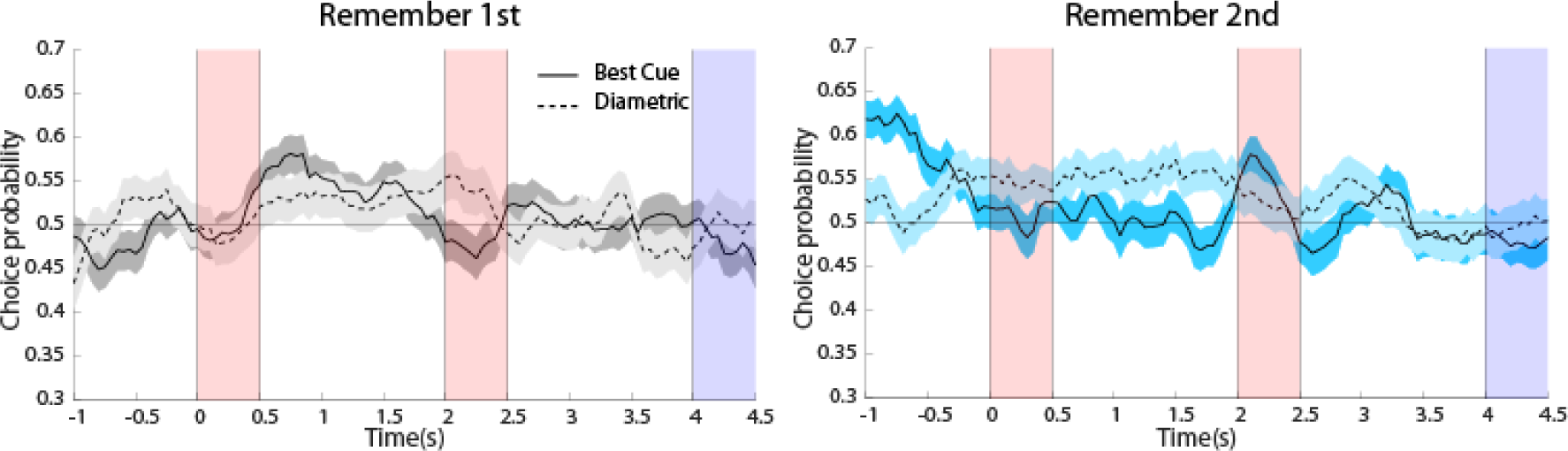
Averaged area under the ROC curve for neurons with significant cue delay activity recorded from PFC (n=75) for remember first task (Left), and remember second task (Right) respectively, plotted as a function of time across the trial. The solid line represents ROC value comparing the distribution of correct and error trials from the preferred cue condition, the dashed line representing the diametric cue condition. The shaded area around represents SEM.

### Population measures in correct and error trials

These results are generally consistent with previous studies that identified changes in firing rate of neurons that generate persistent activity as accounting for behavior in working memory tasks. Recent work has revealed changes in population activity that are not always visible at the level of single neurons or neuron averages, leading to appreciation that the dynamics of population activity can represent cognitive processes and task contexts (11). We therefore employed dimensionality reduction methods to identify population dynamics in the collective activity of neurons and determine how these relate to firing rate changes. Principle Component Analysis (PCA) involving all neurons from the dlPFC and (separately) for the PPC is shown in Fig. 7. Responses are grouped in trajectories with blue or red hue, depending on the location of the cue, at the neurons’ preferred location (blue-hue traces in Fig. 7A), or diametric (red-hue traces in Fig. 7A). The cue was followed by a second stimulus at varying locations (as illustrated in Fig. 1), corresponding to trajectories with slightly different hues. This analysis revealed that activity was relatively stable during the first delay period (gray highlighted rectangle in Fig. 7). Conditions that involved appearance of a stimulus in the neurons’ preferred location were clearly separable from conditions that involved appearance of the stimulus in the diametric location. In the Remember-First task, error trials generated overall similar trajectories; however, the separation of preferred and non-preferred trajectories was now reduced. This difference was evident when we projected delay-period trajectories to the plane defined by the first two principal components (Fig. 8A). The difference in mean distance was significantly reduced (paired t-test, t_99_= 24.2, p<0.001, illustrated in Fig. 8C).

**Figure 7.**
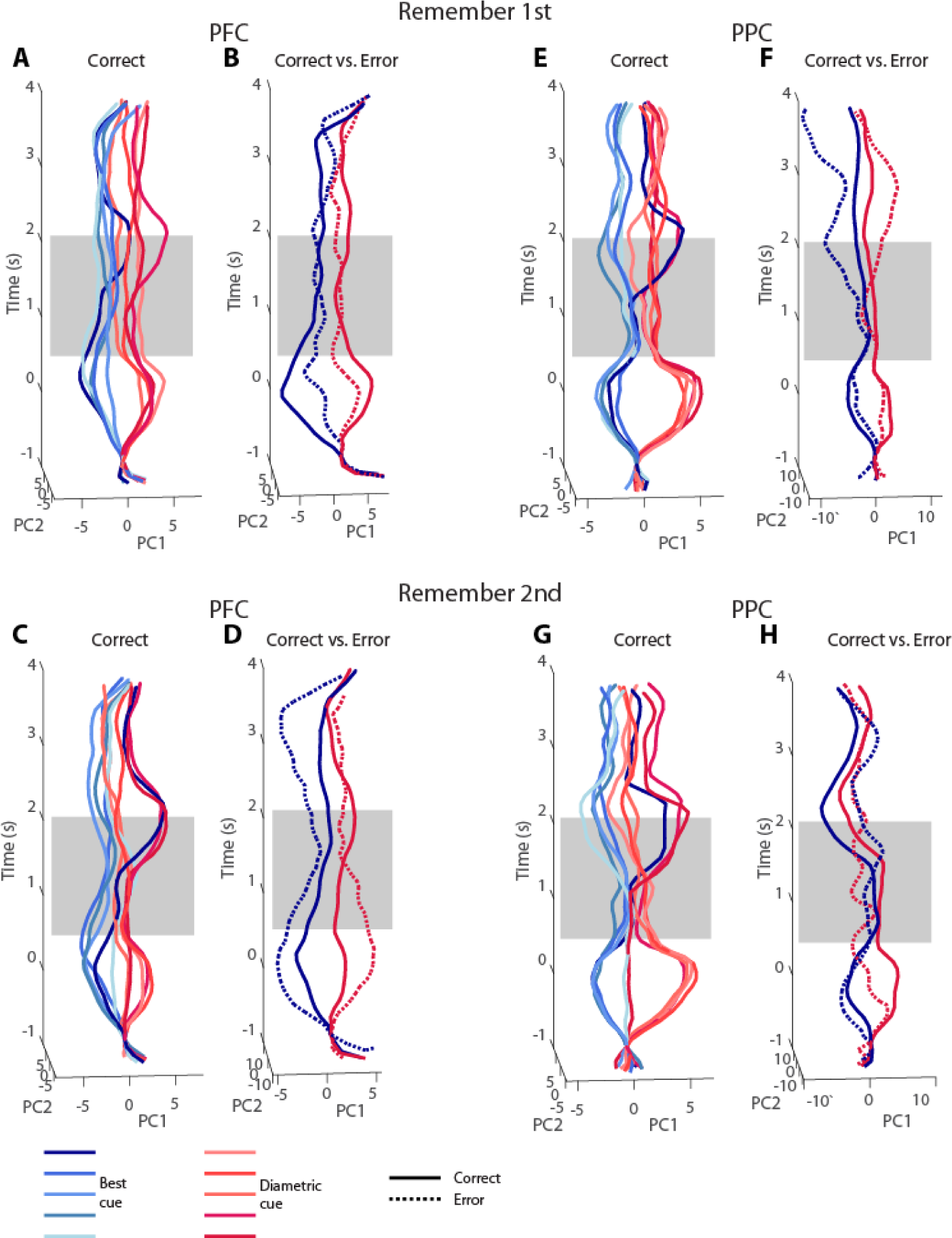
(A) PCA trajectories for correct trials using all neurons recorded from dlPFC (n=205) in the Remember First task. The shaded rectangular area highlights the cue delay period. The blue hue lines indicate the trials with preferred cue condition and red hue lines represent the diametric cue conditioned classes. (B) Averaged correct (solid lines) and error (dotted lines) trial trajectories using all the neurons recorded from dlPFC (n=205) in the remember first task. The shaded rectangular area again highlights the cue delay period. The blue lines indicate the trials with preferred cue condition and red lines represent the diametric cue conditioned classes. (C) As in A, for PFC neurons (n=205) in the remember second task. (D) As in B, for PFC neurons (n=205) in the remember second task (E) As in A, for all PPC neurons (n=241) in the remember first task. (F) Same as in B, for PPC neurons (n=241) in the remember first task. (G) As in C, for PPC neurons (n=241) in the remember second task. (H) Same as in D, for PPC neurons (n=241) in the remember second task.

**Figure 8.**
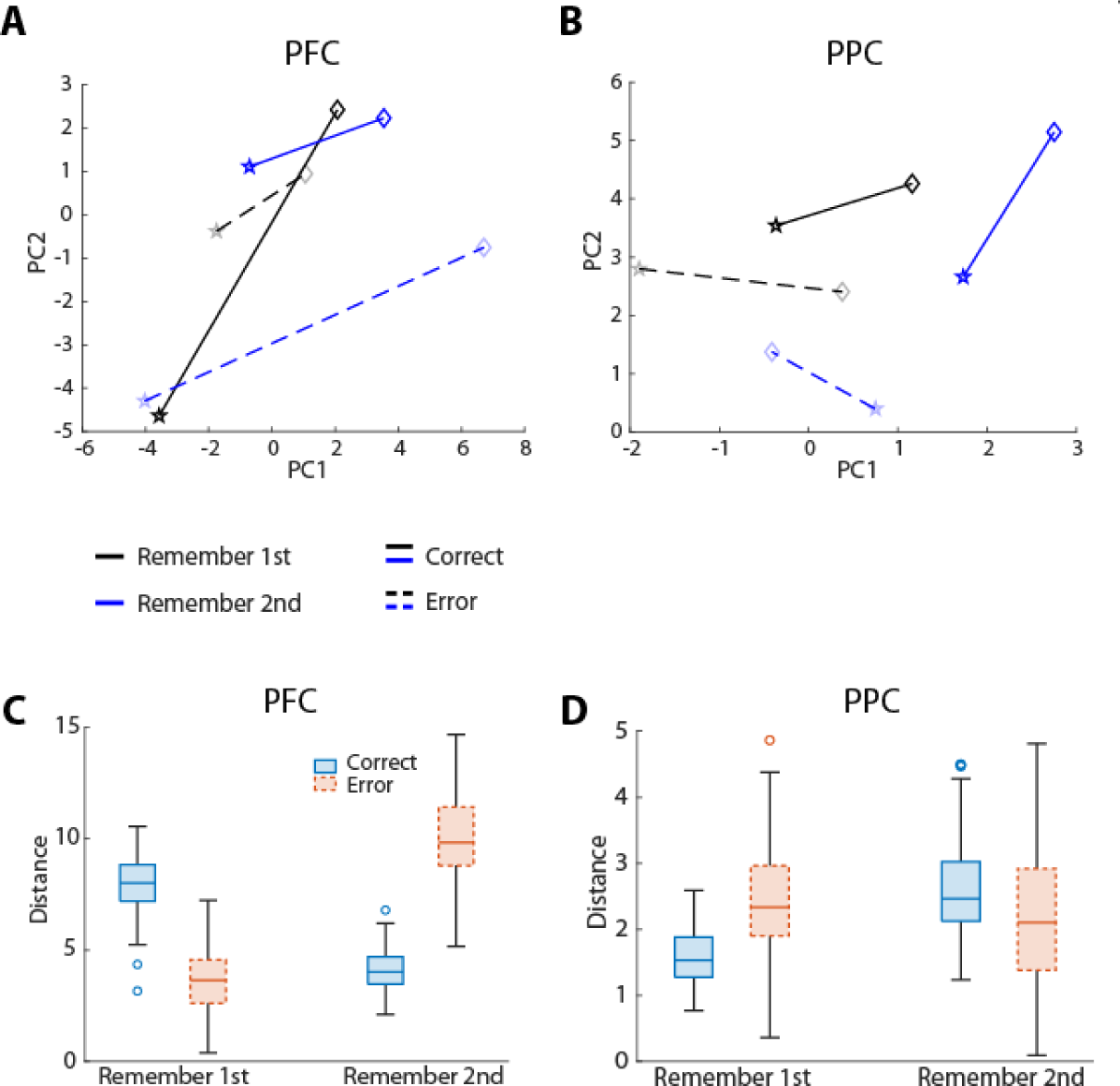
(A) Distance between correct (solid lines) and error (dashed lines) trials when mean points of cue delay trajectories were projected to the state-space plane defined by the first two PCs for PFC neurons (n=205). Here black lines represent the remember first task and blue lines indicate the remember second task. The star shapes indicate the position of the preferred cue and diamond shapes representing diametric cue conditions. (B) Same as in A, for PPC neurons (n=241). (C) Box plots showing distribution of average distances for the delay period (as defined earlier) in correct (blue) and error (red) for PFC neurons (n=205) in both remember first and second task. (D) Same as in C, for PPC neurons (n=241).

Importantly, no such decrease in distance between trajectories of population activity in state space was present for the Remember-Second task (Fig. 7B), and in fact a significant increase was evident (paired t-test, t_99_= −33.2, p<0.001, illustrated in Fig 8A-C). This finding mirrors the change in firing rate discussed earlier; since the first stimulus in this task was to be ignored, increases in firing rate following its presentation were more likely to lead to errors. The opposite trend was present in the PPC for the Remember-First task (Fig. 7C), as the distance between trajectories increased significantly between correct and error trials (paired t-test, t_99_= −8.7, p<0.001, Fig. 8B-D). The distance between stimuli again decreased significantly for error trials in the Remember-Second task (paired t-test, t_99_= 3.13, p=0.002, Fig. 8B-D).

The population analysis produced an additional finding that was not evident in mean firing rates. The axis defined by the best and worst location was shown to rotate between Remember-First and Remember-Second tasks, both for the PFC and PPC (Fig. 8A, 8B). Although the conditions represented by the black and blue traces involved identical stimuli, the axis between best and diametric location appeared at different positions in the principal component plane. This result suggests that different populations of neurons were active in response to stimuli during the delay period in trials of the Remember-First and Remember-Second tasks.

## DISCUSSION

The role of persistent activity in working memory has been reevaluated recently. Although strong evidence links persistent discharges of neurons in the prefrontal cortex with behavioral responses in working memory tasks (21), objections have been raised about its significance (9, 22, 23). At the same time, it has been increasingly appreciated that the collective activity of ensembles of neurons represents more information than is typically evident in the responses of single neurons (11, 12), or average of firing rate across multiple neurons – as persistent activity is typically defined. The results presented here explain how measures of single neuron and population firing are related to working memory behavior.

### Comparison of single-neuron and population measures of activity

Our analysis revealed findings of neuronal firing rate consistent with previous studies, including that persistent activity is generated in the dorsolateral prefrontal and posterior parietal cortex during spatial working memory tasks (24), that activity of neurons that generate persistent activity better predicts working memory recall compared to activity of neurons without persistent activity (8), and that prefrontal neurons better predict performance than posterior parietal neurons, at least in more challenging working memory tasks (25). At the level of mean firing rate, the best predictor of whether a trial might result in an error was whether the delay-period firing of prefrontal neurons generating persistent activity was decreased. Error trials are likely to include lapses, over which firing rate is expected to be generally lower (26). However, the pattern we observed depended specifically on whether the remembered stimulus appeared at the neurons’ preferred location, inside their receptive field. That was the condition best predictive of erroneous responses.

Analysis of the same neuronal datasets using dimensionality reduction methods provided a reflection of these findings at the population level. Population activity was largely captured by a low-dimensional manifold, in which different stimuli were represented at separable loci during working memory, and at a stable subspace even though neuronal activity exhibited considerable dynamics (27). Smaller trajectory deviations of the prefrontal population from their baseline state were more likely to result in errors.

This analysis also revealed changes that were non observable in neuron measures such as average firing rate. Most importantly, we identified rotation of stimulus representations as being associated with different task conditions (whether to remember the first stimulus or the second one). The finding suggests that activation of different ensembles of neurons is needed to maintain stimulus in different working memory tasks. Stimulus rotations have been associated with cognitive operations (28–31) and our findings add to the range of phenomena that suggest that behavior relies on such transformations. Other variables beyond the mean firing rate have also been described that affect behavior, including the variability of neuronal responses from trial to trial (32, 33) and the pair-wise correlation of discharge rates across neurons (34, 35).

### Contributions of prefrontal and posterior parietal cortex in working memory

Neurons in the dorsolateral prefrontal and posterior parietal cortex are co-activated during visuo-spatial working memory tasks, and activity in either area can be predictive of behavior, particularly in simpler tasks that require maintenance of single stimulus in memory, (24, 36). Prefrontal neurons do exhibit some specialization, however; posterior parietal neurons are generally less able to resist distracting stimuli that the subject has to ignore (20, 37), though what is being represented in the activity of each area depends on the specific task a subject is performing (13, 38).

These findings were generally evident in the analysis we performed here. In the posterior parietal cortex, firing rate differences after the presentation of the first stimulus of the remember-first task were generally not well predictive of correct and error trials, precisely because this activity was disrupted by the second stimulus, even though the latter was not relevant for the task. The posterior parietal cortex exhibited the opposite trend compared to the prefrontal cortex in the population analysis. Although the rotation depending on task condition was also evident in posterior parietal populations, the shrinkage or expansion of trajectories moved in the opposite direction than prefrontal ones. These results demonstrate specialization of cortical processes related to working memory and emphasize the role of the prefrontal cortex in the maintenance of working memory (21).

## ACKNOWLEDGMENTS

Research reported in this paper was supported by the NIH National Eye Institute under award numbers R01 EY017077 and P30 EY008126 and by the National Institute of Mental Health under award number R01 MH116675. We wish to acknowledge Chrissy Suell for technical help; Wenhao Dang, Zhengyang Wang, and Rye Jaffe for helpful comments on the manuscript.

